# Sweat and sebum preferences of the human skin microbiota

**DOI:** 10.1101/2022.08.13.503869

**Authors:** Mary Hannah Swaney, Amanda Nelsen, Shelby Sandstrom, Lindsay R Kalan

## Abstract

The microorganisms that inhabit human skin, collectively termed the skin microbiome, must overcome numerous challenges that typically impede microbial growth, including low pH, osmotic pressure, and low nutrient availability. Yet, the skin microbiota thrive on the skin and have adapted to these stressful conditions. Limited skin nutrients are available for microbial use in this unique niche, including those from host-derived sweat, sebum, and corneocytes. Here, we have developed physiologically-relevant, skin-like growth media that is composed of compounds present in human sweat and sebum. We find that skin-associated bacterial species exhibit unique growth profiles in different concentrations of sweat and sebum. The majority of strains evaluated demonstrate a preference for high sweat concentrations, while sebum preference is highly variable, suggesting that the capacity for sebum utilization may be an important driver of skin microbial community structure. Furthermore, these findings provide experimental rationale for why different skin microenvironments harbor distinct microbiome communities. In all, our study further emphasizes the importance of studying microorganisms in an ecologically-relevant context, which is critical for our understanding of their physiology, ecology, and function on the skin.

## INTRODUCTION

Human skin is the primary barrier between the human body and the external environment, carrying out necessary functions that include the prevention of dehydration, regulation of temperature, and protection from mechanical, chemical, and thermal stressors. The skin also has an essential role in protecting against infection by invading pathogenic microorganisms. This protection is conferred in part through the desiccate, nutrient-poor, and slightly acidic nature of the skin, which contributes to a hostile microbial niche that is unwelcoming to pathogen colonization. However, the skin harbors its own community of microorganisms, termed the skin microbiome, that thrives on the skin and has adapted to these seemingly harsh skin conditions.

The skin microbiome, rather than simply occupying the skin niche, carries out a plethora of beneficial functions that range from enhancement of the skin’s physical barrier to early-life education of the immune system (1, 2). A growing list of studies reveal that the skin microbiome contributes to nearly all aspects of skin health and function (3–5). As such, a mechanistic understanding of how the microbiome and its individual members carry out these critical functions is necessary to fully understand the role that the microbiome plays in our health and to develop therapeutics to mitigate skin disorders associated with a disrupted microbiome. For these reasons, the development of model systems to study skin microorganisms in the laboratory under ecologically relevant settings is needed.

In the lab, skin isolates are traditionally grown and studied in rich media that supplies the microorganism with a wealth of nutrients to support rapid and abundant growth. However, nutrient-rich media is not representative of nutrient availability found on the skin, which is limited to minimal host-derived nutrients and metabolites from sweat, sebum, and corneocytes (6). Previous studies have demonstrated that skin commensals utilize nutrients available on the skin, for example through lipase-mediated degradation of sebum lipids by *Cutibacterium acnes* to promote bacterial adherence (7–9), enzymatic cleavage of sphingomyelin to promote barrier integrity and microbial colonization (1), and presumed incorporation of lipids into the bacterial cell membrane (10, 11). In support of investigating skin-associated microbes in an environment that mimics human skin, Horton *et al*. demonstrate that the emerging pathogen and skin colonizer *Candida auris* forms dense multilayer biofilms in synthetic sweat, as compared to routine culture media, providing evidence for how the pathogen persists and spreads in hospital settings (12). Similarly, artificial sweat has been employed in the study of biofilm formation by other clinically-relevant pathogens, including *S. aureus*, and in the culture of mixed skin microbial communities (13, 14). These studies demonstrate that results obtained from laboratory experiments are dependent upon growth conditions, which can vary drastically between the lab setting and the native skin environment.

Across the skin, physiologically distinct skin microenvironments exist as a result of variation in sweat and sebum gland density and distribution, as well as differences in occlusion (exposure to the environment). Skin sites can broadly be classified as sebaceous, moist, dry, or foot microenvironments. Sebaceous sites (face, back) are characterized by a higher density of hair follicles and sebaceous glands, which secrete lipid-rich sebum (15). Moist sites (belly button, bend of the elbow) are often semi-occluded and can have more abundant and active sweat glands, which lead to increased moisture levels (16, 17). Dry sites (forearm, back of the hand) have a lower density of sweat and sebaceous glands (15, 16). Lastly, foot sites (toe web space, heel) are generally occluded (as a result of shoe wearing) with increased moisture levels and have a lower temperature than the majority of other skin sites (16, 18). Numerous studies have demonstrated that the microbial communities of these distinct microenvironments are unique in their structure and function (19–22), suggesting that the observed community differences between microenvironments are in part due to differential microbial utilization of available nutrients and/or microbial inhibition by human skin secretions.

To more accurately mimic the native skin environment in the laboratory, we developed a custom human skin secretion media for cultivating skin commensals that includes physiologically-revelant compounds found in both sweat and sebum. Because the microbial communities across the biogeography of the skin are driven by distinct microenvironments, we hypothesize that skin-associated bacteria isolated from these microenvironments will exhibit differential growth in sweat and sebum. Using 15 phylogenetically-diverse bacterial skin strains isolated from healthy skin and the skin pathogen *Staphylococcus aureus*, we show that each strain has a unique sweat and sebum preference profile. We find that overall, skin-associated bacterial species have a strong preference for high sweat concentrations. In contrast, sebum preference varies, with strains exhibiting a range of lipophilic to lipophobic phenotypes. Overall, our findings provide additional evidence to support the hypothesis that skin microbial community structure varies across skin microenvironments as a result of differences in skin secretions and nutrient availability. Understanding microbial nutrient preferences in physiologically relevant conditions is a critical step towards developing model skin microbiome systems and interrogating microbial interactions.

## RESULTS

To systematically study skin commensal growth preferences in nutrients representative of those found on the skin, we developed an artificial sweat and sebum that can be supplemented to minimal medium in a dose dependent manner. We developed the minimal medium (termed here as basal medium) to provide minimal nutrients required to support growth of most skin-associated bacteria. The composition consists of salts, glucose as a carbon source, and select amino acids and vitamins (Supplemental Table 1).

### Artificial sweat development

Human eccrine sweat is a complex mixture of solutes that is secreted from eccrine sweat glands, which are the most numerous sweat gland found across the body surface area and are responsible for the highest volume of sweat production (23). Eccrine sweat, composed of mainly salt water, contains a variety of other solutes in varying concentrations, including lactate, urea, glucose, bicarbonate, amino acids, electrolytes (in addition to Na^+^ and Cl^-^), and vitamins (23, 24). To develop an artificial sweat that is consistent with the composition of human eccrine sweat and is suitable for microbial growth, we developed a formulation considering previously described synthetic sweat media (12, 14) and through extrapolating eccrine sweat constituents and their concentrations as reported in the literature (Supplemental Material S1). The composition of the artificial sweat used in this study is listed in Table 1.

**Table 1.** Artificial sweat composition.

| Artificial Sweat Component | Composition (mg) |
| --- | --- |
| Salts |  |
| Sodium Chloride (g) | 2 (g) |
| Sodium Pyruvate | 50 |
| Monosodium Phosphate anhydrous | 10 |
| Calcium Sulfate dihydrate | 50 |
| Potassium H Carbonate | 170 |
| Carbon Sources |  |
| D-Glucose | 20 |
| Amino Acids |  |
| Glycine | 50 |
| L-Leucine | 25 |
| L-Serine | 80 |
| L-Alanine | 40 |
| L-Arginine | 50 |
| L-Histidine | 100 |
| L-Valine | 20 |
| L-Isoleucine | 15 |
| L-Lysine HCl | 35 |
| L-Phenylalanine | 15 |
| L-Tyrosine | 20 |
| L-Glutamine | 5 |
| L-Aspartic Acid | 25 |
| L-Glutamic acid | 35 |
| L-Proline | 8 |
| L-Ornithine HCl | 50 |
| L-Citrulline | 50 |
| L-Tryptophan | 20 |
| L-Asparagine | 10 |
| L-Threonine | 35 |
| Other |  |
| Creatinine | 5 |
| Lactic Acid (88%) (g) | 0.5 (g) |
| Urea (g) | 0.5 (g) |

Values are expressed in milligrams (per 1 liter of medium) unless noted otherwise

### Artificial sebum development

Human sebum is a complex mixture of lipids that is secreted from sebaceous glands, which are distributed across the body and most densely located on the scalp, forehead, and face (25). Sebum is composed of triglycerides, free fatty acids, squalene, wax esters, cholesterol esters, and cholesterol (26). To provide a sebum-like lipid source when culturing skin commensals, we developed an artificial sebum similar in composition to sebum L, which was developed by Lu *et al*. and found to be physiochemically consistent with human sebum (27). The composition of the artificial sebum used in this study is listed in Table 2. Because this lipid formulation is not miscible with water, we found that a ratio of one part artificial sebum to three parts Tween 80 allowed for solubilization of the artificial sebum in aqueous solution. Tween 80 is a non-ionic surfactant that contains oleic acid (an abundant fatty acid present on the skin) and is commonly used in the culture of skin microorganisms (28, 29).

**Table 2.** Artificial sebum composition.

| Artificial Sebum Component | Composition (%w/w) |
| --- | --- |
| Squalene | 15 |
| Cholesterol | 1.2 |
| Cholesterol Oleate | 2.4 |
| Waxes |  |
| Paraffin Wax | 10 |
| Hexadecyl Palmitate | 15 |
| Fatty Acids |  |
| Oleic Acid | 1.4 |
| Myristic Acid | 2.5 |
| Lauric Acid | 2.5 |
| Palmitic Acid | 5 |
| Triglycerides |  |
| Olive Oil | 10 |
| Coconut Oil | 10 |
| Cottonseed Oil | 25 |

### Selection of bacterial isolates from diverse taxa and numerous skin sites

To evaluate the growth of skin commensals in media containing the artificial sweat and sebum, we selected 15 phylogenetically diverse bacterial skin strains isolated from healthy skin in our laboratory, as well as the skin pathogen *S. aureus*, for testing. These skin isolates represent species from the four most abundant phyla present in the skin microbiome: Actinomycetota (n=10 strains), Bacteroidota (n=1 strain), Pseudomonadota (n=2 strains), and Bacillota (n=3 strains) (Figure 1A, Supplemental Material S2). Furthermore, this collection, which contains strains isolated from 8 distinct skin sites (alar crease, occiput, nare, umbilicus, antecubital fossa, axillary/groin, volar forearm, and toe web space), includes representatives from the 4 skin microenvironments: sebaceous (n=2 strains), moist (n=8 strains), dry (n=3 strains), foot (n=2 strains) (Figure 1B and 1C). We have observed that less abundant or rare species are more frequently isolated from moist skin sites, therefore we selected numerous strains isolated from this microenvironment to increase the phylogenetic diversity of the isolates tested.

**Figure 1.**
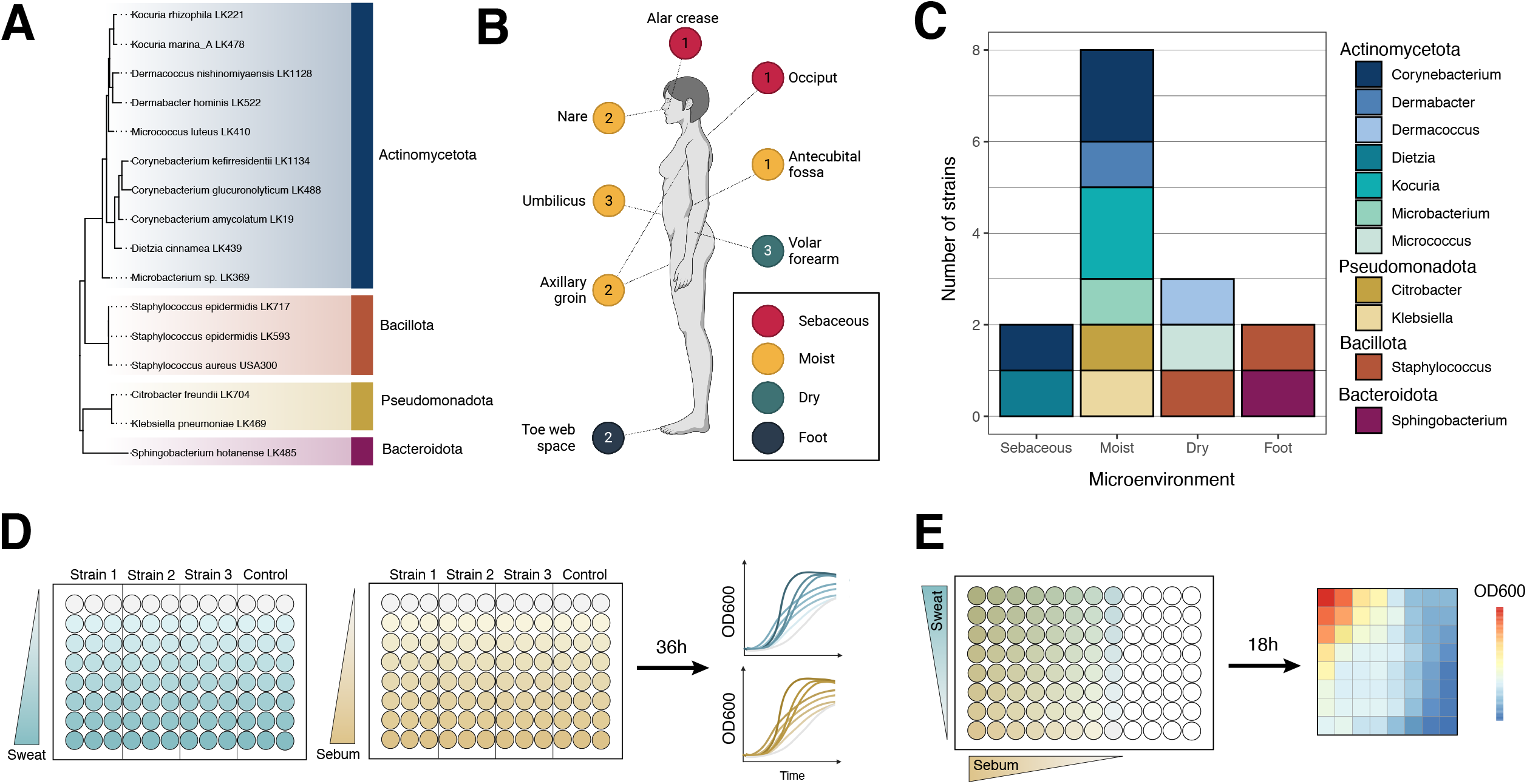
Overview of skin strains and experiment design. A) 15 strains isolated from healthy subjects and *Staphylococcus aureus* USA300 were selected for use in this study and include representatives from the Actinomycetota (blue), Bacillota (orange), Pseudomonadota (yellow), and Bacteroidota (dark pink) phyla. 16S rRNA sequences from each strain were aligned, and an approximately-maximum-likelihood phylogenetic tree was inferred. The tree was rooted using archaeum *Halobacterium salinarum*. B) The skin strains were isolated from 8 skin sites, with the number of strains isolated from each site indicated. The axillary groin site is a combined swab of the axilla and groin skin. Colored circles represent skin microenvironment category: sebaceous (red), moist (yellow), dry (green), foot (blue). C) The number of strains isolated from each microenvironment and their respective genus taxonomic assignment. D) Experimental overview of the growth curve assays. Per one 96-well plate, three strains were inoculated into increasing concentrations of artificial sweat or sebum media and incubated at 37ºC for 36 hours. OD_600_ measurements were taken at even intervals. E) Experimental overview of the checkerboard-like assay. Per one 96-well plate, one strain (representing a single biological replicate) was inoculated into increasing concentrations of combined artificial sweat and sebum media. The plate was incubated at 37ºC for 18 hours, at which point an endpoint OD_600_ measurement was taken.

### Skin strains exhibit preference for high sweat concentrations

The 16 selected strains were then cultured for 36 hours in basal medium containing increasing concentrations of sweat for growth curve analysis (Figure 1D). Sweat concentrations were tested from 0X (only basal medium) through 4X (only artificial sweat, no basal medium). We selected concentrations above 1X artificial sweat in order to test the tolerance of the skin strains at higher salt and solute concentrations and to account for variation in sweat composition across individuals. Furthermore, these higher sweat concentrations may be physiologically relevant, as skin surface sweat rapidly evaporates and may leave residual sweat constituents, which has been observed previously during sweat collection (30–32).

We observed that generally, the skin strains tested either exhibited similar growth patterns at all sweat concentrations tested, apart from 4X sweat, or exhibited increased growth with increasing sweat concentration (Figure 2A). This concentration-dependent increase in growth is particularly evident for the two *Staphylococcus epidermidis* strains, which grew very poorly and variably in basal medium alone or in low concentrations of sweat. Furthermore, we observed minimal or no growth over time when strains were grown in 4X sweat, which contained no basal medium, suggesting either that certain basal medium components are required for rapid growth or that the concentration of sweat solutes was too high or not tolerated well for the strains tested. To quantify the growth curves for each strain and sweat concentration tested, we calculated maximum growth rate and area under the curve (AUC), which integrates information about lag phase, growth rate, and carrying capacity (33). We observed either little AUC change across sweat concentrations or a concentration-dependent increase for these two metrics (Figure 2B, Supplemental Figure 1A). We confirmed this finding by performing a linear regression with the AUC data to assess sweat preference of each strain, which revealed that most strains either exhibit no preference for sweat or a high concentration preference (Figure 2C).

**Figure 2.**
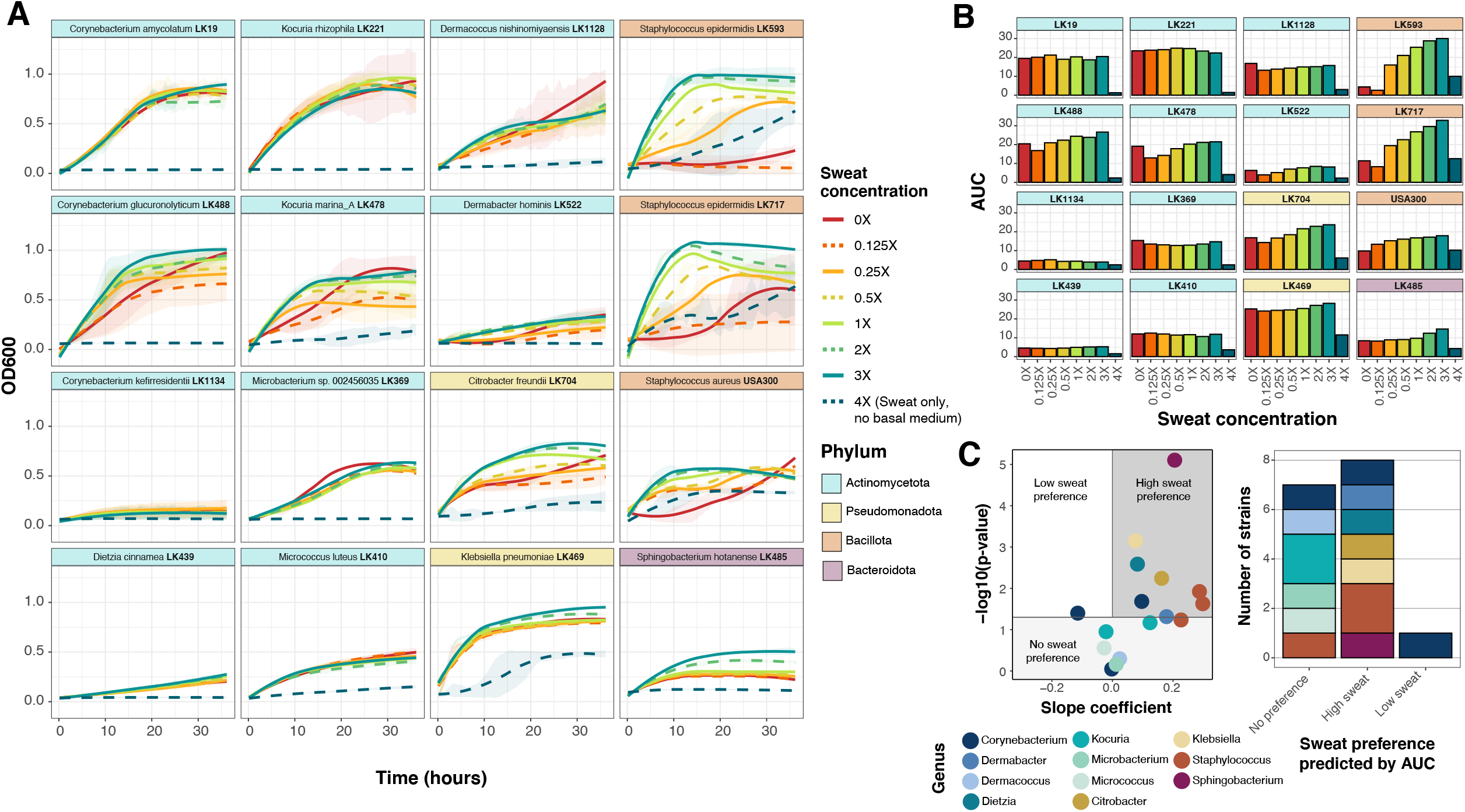
Skin species exhibit concentration-dependent growth in artificial sweat. A) For each strain, growth curves were collected in eight concentrations of artificial sweat media over 36 hours. Strains are ordered by phylum-level taxonomic assignment. 0X sweat contains only basal medium, and 4X sweat contains only artificial sweat. Curves were generated using loess smoothing for averaged OD_600_ values from at least 2 biological replicates with 3 technical replicates each. Ribbons represent the standard deviation across biological replicates. B) Area under the curve (AUC) was calculated for each strain and each artificial sweat concentration. Strain order is identical as in panel A. C) Linear regression was performed with min-max normalized AUC data. For each strain, slope coefficient (β) and p-value for the model are plotted. Strains are classified into sweat preference based on the following coefficient and p-value cutoffs: low sweat = β < 0 and p-value < 0.05; high sweat = β > 0 and p-value < 0.05; no sweat preference = p-value > 0.05.

We also tested pathogen *S. aureus* in the sweat media, which revealed a sweat concentration-dependent increase in growth, similar to the *S. epidermidis* strains. However, unlike these commensal staphylococci, growth was comparatively much less robust in higher sweat concentrations, which supports the notion that skin commensals are highly adapted to the skin environment and available nutrients. Indeed, Staphylococci have numerous osmoprotectant transport systems for providing resistance to changing osmolarity (34), which is likely exhibited on the skin as a result of the production and evaporation of sweat. It has been suggested that *S. epidermidis* may have increased presence or expression of these protective systems compared to *S. aureus* (35).

### Skin strains exhibit a range of sebum concentration preferences

To evaluate growth of the skin strains in artificial sebum media, sebum concentrations were tested from 0% (only basal medium) through 0.25%. We selected 0.25% as the maximum concentration because above this, the lipids reach their solubility limit. We found that growth curve patterns with differing sebum concentrations varied drastically across strains. For example, certain strains including *Corynebacterium kefirresidentii* and *Dietzia cinnamea* exhibit a concentration-dependent increase in growth with increasing sebum concentration, whereas other strains such as *Kocuria rhizophila* and *Microbacterium* sp. LK369 exhibit a concentration-dependent decrease in growth (Figure 3A). Furthermore, some strains including *Micrococcus luteus* and *Dermacoccus nishinomiyaensis* exhibited relatively similar growth curves regardless of sebum concentration. These patterns are similarly reflected in the AUC and max growth rate for each strain and sebum concentration (Figure 3B, Supplemental Figure 1B). We then classified the isolates by the sebum preference, determined by linear regression of AUC data, into the following categories: low sebum, high sebum, or no sebum preference. We found that the skin strains demonstrate a range of sebum preferences, with multiple strains falling into each category (Figure 3C).

**Figure 3.**
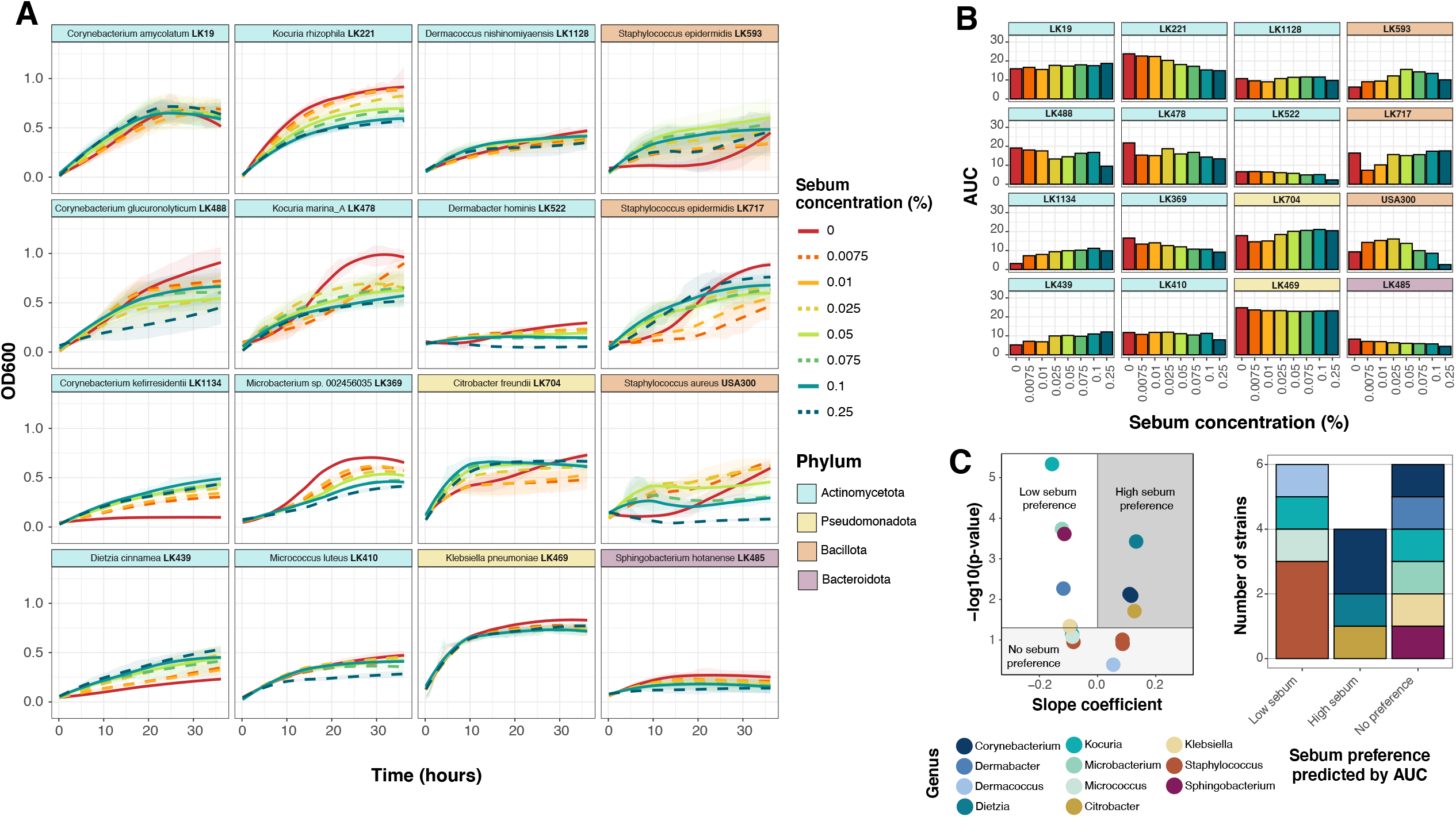
Growth in artificial sebum varies by skin species. A) For each strain, growth curves were recorded in eight concentrations of artificial sebum media over 36 hours. Strains are ordered by phylum-level taxonomic assignment. 0% sebum contains only basal medium. Curves were generated using loess smoothing for averaged OD_600_ values from at least 2 biological replicates with 3 technical replicates each. Ribbons represent the standard deviation across biological replicates. B) Area under the curve (AUC) was calculated for each strain and each artificial sebum concentration. Strain order is identical as in panel A. C) Linear regression was performed with min-max normalized AUC data. For each strain, slope coefficient (β) and p-value for the model are plotted. Strains are classified into sebum preference based on the following coefficient and p-value cutoffs: low sebum = β < 0 and p-value < 0.05; high sebum = β > 0 and p-value < 0.05; no sebum preference = p-value > 0.

### Growth profiles of skin strains in sweat and sebum cluster by phylum

To assess growth of the skin strains in media containing both artificial sweat and artificial sebum, we developed a checkerboard-like assay for testing strain growth in 8 concentrations of sebum (0% through 0.125%) across 8 concentrations of sweat (0X through 2X) (Figure 1E). Strains were cultured for 18 hours, after which an end point OD_600_ measurement was collected. We selected this time point because growth curve data suggests that 18 hours is when strains had reached early stationary phase and exhibited the largest differences in OD_600_ among the varying sweat or sebum concentrations (Fig 2A & 3A). Checkerboard assay results demonstrate a wide range of growth phenotypes across the different sweat and sebum concentrations for each strain (Figure 4, Supplemental Figure 2). In certain cases, highly similar growth patterns occur for phylogenetically related species (e.g., *Citrobacter freundii* and *K. pneumoniae* from the Pseudomonadota phylum, or the three strains from the *Staphylococcus* genus). However, across strains from the Actinomycetota phylum, a range of growth patterns and sweat/sebum preferences exist. Even within a single genus, as for *Corynebacterium* and *Kocuria* for which multiple species representatives are included, multiple phenotypes are demonstrated.

**Figure 4.**
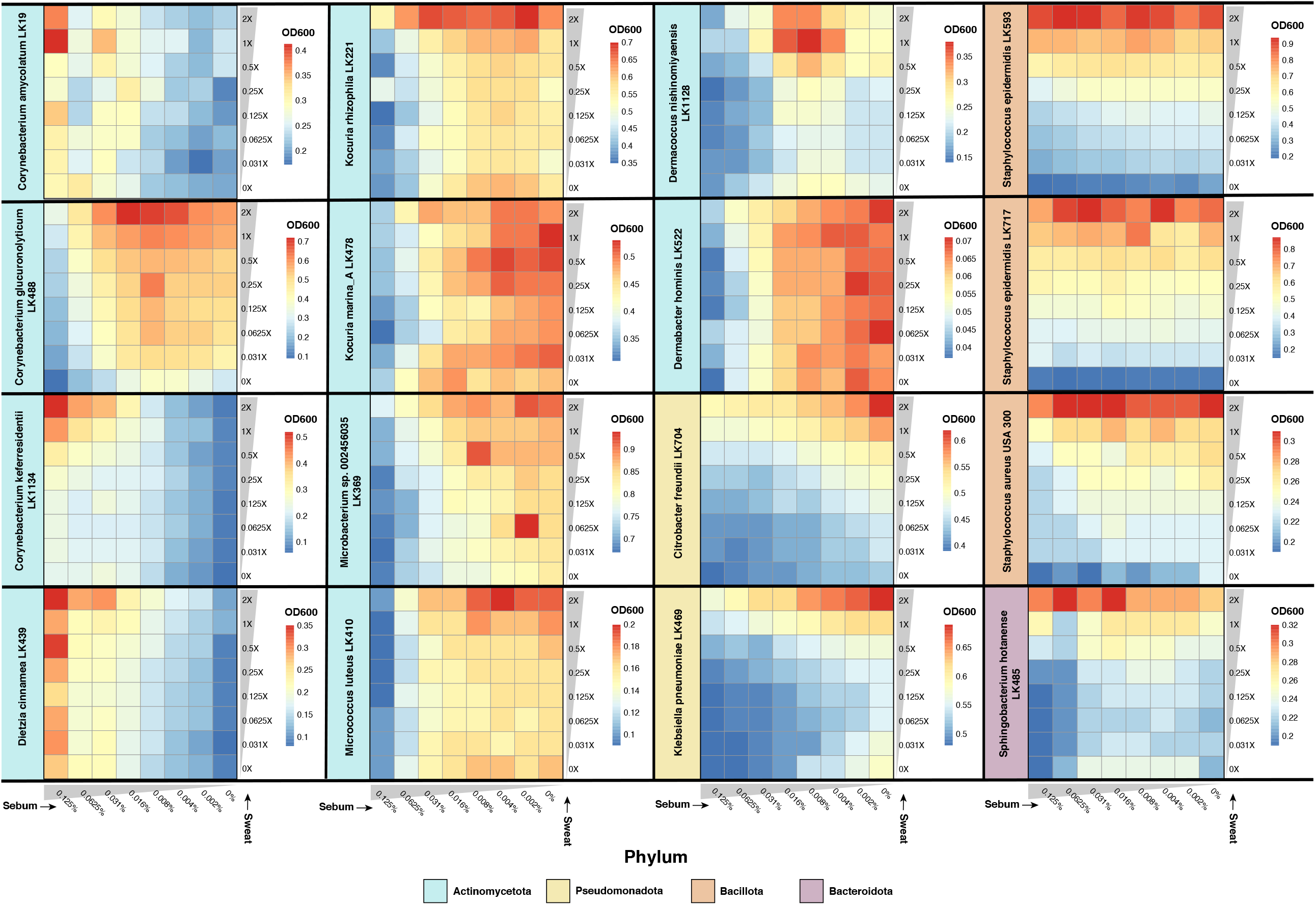
Sweat and sebum checkerboard assays reveal species-specific growth patterns. Strains were cultured in unique combinations of artificial sweat media (0, 0.031, 0.0625, 0.125, 0.25, 0.5, 1, 2X) and artificial sebum media (0, 0.002, 0.004, 0.008, 0.016, 0.031, 0.0625, 0.125%) for 18 hours, after which an endpoint OD_600_ reading was recorded. At least three replicates were collected for each strain, and OD_600_ values from uninoculated media were subtracted from experimental OD_600_ values. Average OD_600_ values of the replicates are shown.

To group together strains by growth profile similarity, we employed hierarchical clustering of the checkerboard data, which grouped strains with similar profiles into three main clusters. We further observed clustering by phyla (Figure 5A). This phylogenetic signal was supported by a tanglegram analysis, which demonstrated correspondence between the checkerboard dendrogram and the 16S rRNA phylogeny for the Bacillota and Pseudomonadota subtrees. Within the Actinomycetota phylum, two clusters formed, with one cluster consisting of strains that exhibited increased growth with increasing sebum concentrations (e.g., *Cornyebacterium amycolatum*), and the other cluster consisting of strains that exhibited decreased growth with increasing sebum concentrations (e.g., *Kocuria marina*). Consistent with this analysis, we performed principal components analysis (PCA) and identified three distinct groupings, with two groups consisting of strains from the Actinomycetota phylum and the other consisting of strains from the Bacillota, Pseudomonadota, and Bacteroidota phyla (Figure 5B). When considering skin microenvironment, strong grouping of strains by the microenvironment they were isolated from was not observed, with the exception of sebaceous site-derived isolates *C. kefirresidentii* and *Dietzia cinnamea*.

**Figure 5.**
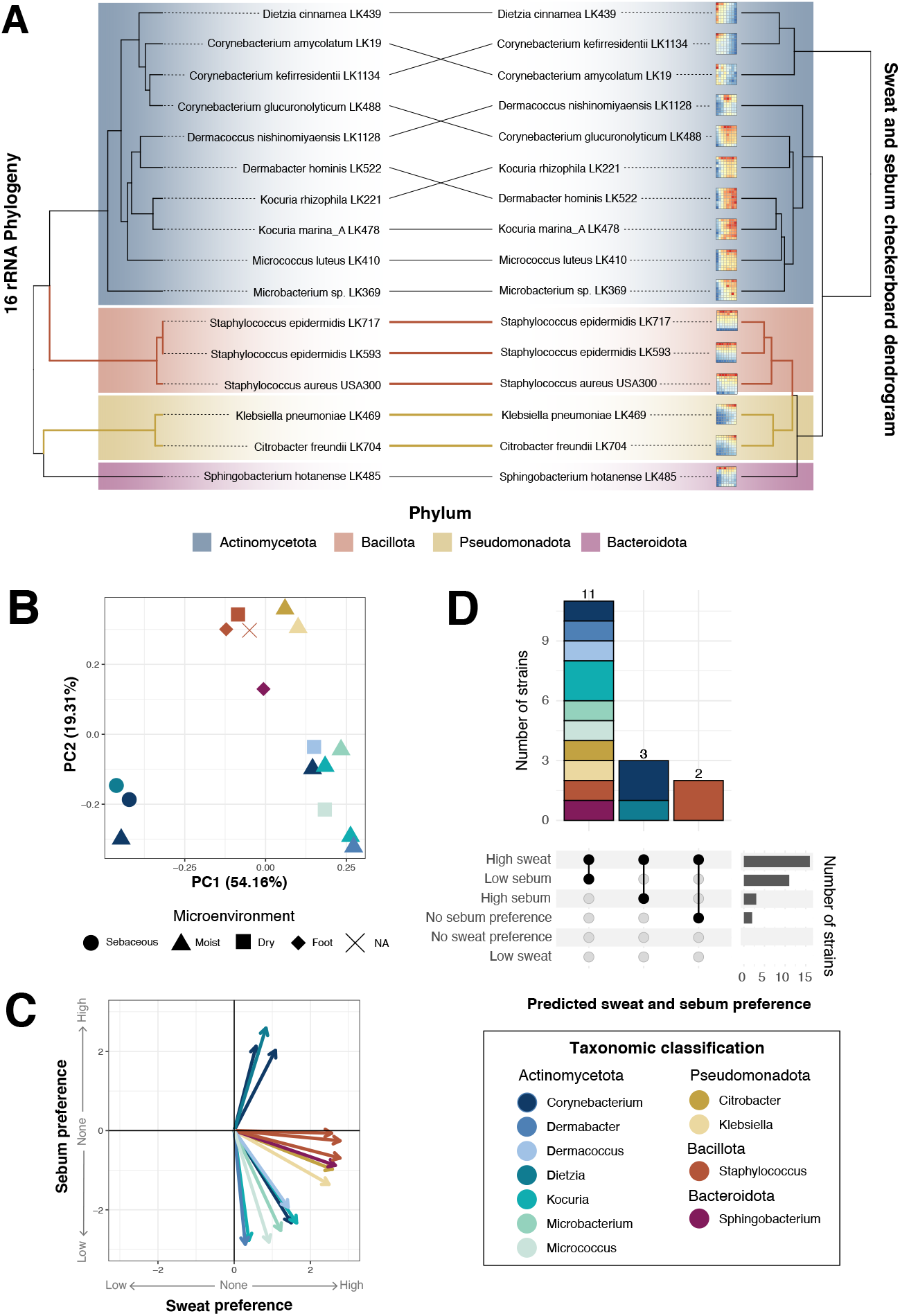
Skin species group by sweat and sebum preference. A) Hierarchical clustering was performed on z-score normalized data from the sweat and sebum checkerboard assays. Results are plotted as a tanglegram with the resulting dendrogram (right) and the 16S rRNA phylogeny from Figure 1A (left). B) Principal components analysis and C) multiple linear regression were performed with the z-score normalized data. In C), the slope coefficients for sweat concentration and sebum concentration from the regression analysis are plotted as an (x,y) coordinate, respectively. D) Strains are classified by sweat and sebum preference based on the following slope coefficient (β_Sweat_ or β_Sebum_)and p-value cutoffs: low sweat = β_Sweat_ < 0 and *p*-value < 0.05; high sweat= β_Sweat_ > 0 and p-value < 0.05; no sweat preference = p-value > 0; low sebum = β_Sebum_ < 0 and p-value < 0.05; high sebum = β_Sebum_ > 0 and p-value < 0.05; no sebum preference = p-value > 0. Strains are colored by phylum (A) or genus (B-D).

### Skin strains prefer high sweat concentration but varied sebum concentration

To quantify the strength of sweat and sebum preference for each strain, we performed a multiple linear regression (MLR) analysis of the checkerboard data and ordination of the the slope coefficients, which represent the rate of change of OD_600_ with increasing sweat or sebum concentration. This revealed that most strains have a preference towards higher sweat concentrations, with *S. epidermidis* LK717, *S. epidermidis* LK593, *S. aureus, Sphingobacterium hotenanse*, and *Citrobacter freundii* demonstrating the strongest preference for high sweat concentrations (Figure 5C). For these strains, sweat concentration had a relative importance of 99.94%, 99.15%, 94.15%, 90.17%, and 87.48%, respectively, in the the regression model (Supplemental Material S3). However, certain strains including *Dermabacter hominis* and *Kocuria marina* appear to have relatively minimal preference for sweat concentration, with low relative importance of 1.02% and 1.91%. On the other hand, sebum preference varied drastically for each strain, with only *C. amycolatum, D. cinnamea*, and *C. kefirresidentii* exhibiting a preference for high sebum concentrations (relative importance of 93.24%, 90.58%, and 77.78%). Both *S. epidermidis* strains exhibited little preference for any sebum concentration (relative importance of 0.06% for LK717 and 0.85% for LK593). The remaining strains demonstrated a preference for low sebum concentrations, with *D. hominis, K. marina*, and *M. luteus* showing the strongest preference for low sebum and relative importance of 98.98%, 98.09%, 90.47%, respectively.

Although we did not observe grouping based on skin microenvironment, we were interested if skin microenvironment where the strains were isolated was reflected in the strength of sweat and sebum preference. We observed that the two strains isolated from the foot microenvironment, *S. epidermidis* LK717 and *S. hotanense*, exhibited strong preference for high sweat concentrations (Supplemental Figure 3). Furthermore, the two skin strains isolated from sebaceous skin sites, *C. kefirresidentii* and *D. cinnamea*, were 2 of 3 strains out of the total 16 tested that showed preference for high sebum concentrations. We did not observe a pattern for the other isolates from dry or moist microenvironments.

The strains were then classified by their sweat and sebum preference according to MLR slope coefficients and corresponding p-values. This revealed that 11 of the 16 strains exhibit high sweat and low sebum preference, 3 strains exhibit high sweat and high sebum preference, and 2 strains exhibit high sweat preference but no sebum preference (Figure 5D). Overall, a range of sweat and sebum preferences are exhibited by the skin strains, with preference for high sweat concentration most common.

## DISCUSSION

The study of microorganisms in conditions similar to their native environment is crucial for characterizing microbial physiology, ecology, and function. Particularly, the skin microbiota have adapted to the desiccate, nutrient-limited conditions on the skin, and as such are equipped to metabolize the available nutrients derived from sweat, sebum, and corneocytes present in the skin environment. Yet, skin microorganisms are often supplemented with a surplus of nutrients when they are cultured in nutrient-rich media in the lab. In this study, we developed defined artificial sweat and sebum media that supports growth of diverse skin bacteria from the four most abundant phyla present on the skin. These artificial formulations contain physiologically-relevant compounds that are found in human sweat and sebum and are based upon numerous studies that have characterized the composition of human sweat and sebum (18, 24, 27, 36–45), as well as previously developed artificial formulations for the growth of microorganisms (12, 14).

Using the developed artificial sweat and sebum, we systematically tested the growth of 15 bacterial strains isolated from healthy human skin as well as the skin pathogen *S. aureus*. We found that the strains overall have a strong preference for media with high sweat concentrations and generally exhibit more growth when supplemented with artificial sweat. Eccrine sweat glands, which produce a salty secretion that our artificial sweat mimics, are distributed across almost all of the human body (23). Therefore, it could be expected that the skin microbiota have adapted to utilize this ubiquitous source of nutrients for their metabolism and growth. While to our knowledge no prior studies have systematically evaluated individual skin commensal growth dynamics in sweat-like conditions, Nagarajan and colleagues cultured *Staphylococcus, Cutibacterium*, and *Micrococcus* skin isolates in filter-sterilized sweat collected from human subjects to reveal that *Staphylococcus* species metabolized sweat to produce malodor-associated compounds (46). Furthermore, Callewaert *et al*. were able to successfully use an artificial sweat to culture skin microbial communities for up to 21 days which closely resembled the native microbial community from which they were sampled (14). Thus, our study builds upon previous knowledge to provide evidence that the skin microbiota utilize sweat as a nutrient source for metabolic function and growth.

In particular, we observed that the growth of *S. epidermidis*, and to a lesser extent *S. aureus*, was highly responsive to increasing sweat concentrations, suggesting that some constituent of the artificial sweat could be required or preferred for abundant growth. However, the exact component(s) of the artificial sweat that results in this concentration-dependent growth increase is unknown. It is predicted that *Staphylococcus* and *Corynebacterium*, and likely the other skin microbiota, utilize urea and amino acids in sweat as a nitrogen source, and glucose, lactic acid, and amino acids as carbon sources (18, 47). Human sweat itself is considered to be hypotonic (∼0.2% NaCl), but the skin surface is likely much higher in salt content as a result of residual salt deposits after sweat evaporation. While the highest NaCl concentration tested in this study only reached 1.05% (4X artificial sweat) and is not considered a high salt concentration, many skin microorganisms are highly halotolerant and capable of withstanding drastic changes in osmotic pressure, including *Staphylococcus* and *Corynebacterium* species (47–53). In addition, nearly all other genera in this study have reports of species being identified as halotolerant (54–59).

In contrast to the observed preference for higher concentrations of artificial sweat for the skin strains tested, we found that strain preference for artificial sebum concentration varies, even among strains within the same genus. For example, certain strains, such as those from the *Corynebacterium* and *Dietzia* genera, exhibited lipophilic tendencies and showed a strong preference or even requirement for higher sebum concentrations. This is likely a result of a lipid auxotrophy, which is exhibited by certain microorganisms colonizing the skin, including several *Corynebacterium* species and fungal species within the *Malassezia* genus (6). For example, the human skin commensal *Corynebacterium jeikeium* lacks a fatty acid synthase and is thus dependent on exogenous fatty acids. Fatty acids are the building blocks for cornyomycolic acids, the characteristic major constituents of the cell envelope for most *Corynebacterium* species (60), and are thus necessary for Corynebacterial growth. Indeed, we observed that *C. kefirresidentii*, one of the most abundant species colonizing the skin (61), requires sebum for growth, suggesting that in minimal media, *C. kefirresidentii* is a lipid auxotroph.

While some of the tested skin strains preferred sebum, many strains demonstrated inhibited growth with increased sebum concentration. Sebum itself, and particularly the free fatty acids that make up a portion of this lipid-rich secretion, have antimicrobial properties, which could explain why many skin strains in this study have a low tolerance for sebum (29, 62, 63). Interestingly, over 80% of *S. epidermidis* strains can esterify fatty acids to cholesterol, which may protect them from sebum’s antimicrobial effects (47, 64). In the checkerboard assay, regardless of sebum concentration, both *S. epidermidis* strains demonstrated similar growth patterns, and sebum concentration had very little relative importance in the linear model for describing growth with increasing sweat and sebum concentration. Therefore, certain skin commensals, including *Staphylococcus* species, may be well equipped to tolerate the antimicrobial properties of sebum on the skin.

The different microenvironments across the skin harbor distinct skin microbial communities (19). Presumably, these unique communities exist in part because of the physiological differences of the skin, including sweat gland, sebaceous gland, and hair follicle density and distribution, which lead to differences in sweat and sebum secretion. Here, we demonstrate that skin isolates exhibit unique growth profiles in sweat and sebum and have specific sweat and sebum concentration preferences, providing *in vitro* evidence to support the hypothesis that microbial community organization differs across skin microenvironments as a result of differences in skin secretions and nutrient availability. Our results further suggest that sebum may be a dominant driver of these differences, as sebum preference differed drastically from strain to strain. Indeed, sebaceous skin sites are dominated by species that can thrive in this lipid-rich, antimicrobial environment. For example, the anaerobic, aerotolerant *Cutibacterium acnes* metabolizes and breaks down sebum components to supports its own growth (65, 66) and to control growth of other microbes (67), contributing to its success at colonizing sebaceous skin sites.

A limitation of this study is that only skin bacterial strains that grow aerobically were selected for testing, therefore future studies to interrogate how anaerobes utilize sweat and sebum is necessary. Furthermore, we did not assess chemical stability of our artificial sweat and sebum formulations. However, we have observed consistent results using the media stored at 4ºC and protected from light over a period of approximately two months. Lastly, antimicrobial peptides produced by the host are present in skin secretions and undoubtedly influence microbial growth (68–70), therefore our findings do not reflect the effect that these important molecules may also have on commensal growth and microbiome structure.

In conclusion, we have developed artificial sweat and sebum media that supports the growth of a wide range of skin microorganisms. We also show that skin strains demonstrate unique growth profiles in different concentrations of sweat and sebum and have distinct preferences for sweat and sebum concentration. These findings provide evidence to support why different skin microenvironments, characterized by their physiological properties, harbor unique microbial communities. Importantly, our findings support the use of physiologically-relevant media in the study of skin microorganisms. Future studies using metabolomics to characterize metabolic output of skin commensal growth in the artificial sweat and sebum media and transcriptomics to examine pathways that may be involved in sweat and sebum utilization will provide important insight into how the skin microbiota have adapted to thrive on the skin, and how we can harness these adaptations in the context of health and disease.

## FIGURE LEGENDS

**Supplemental Figure 1.**
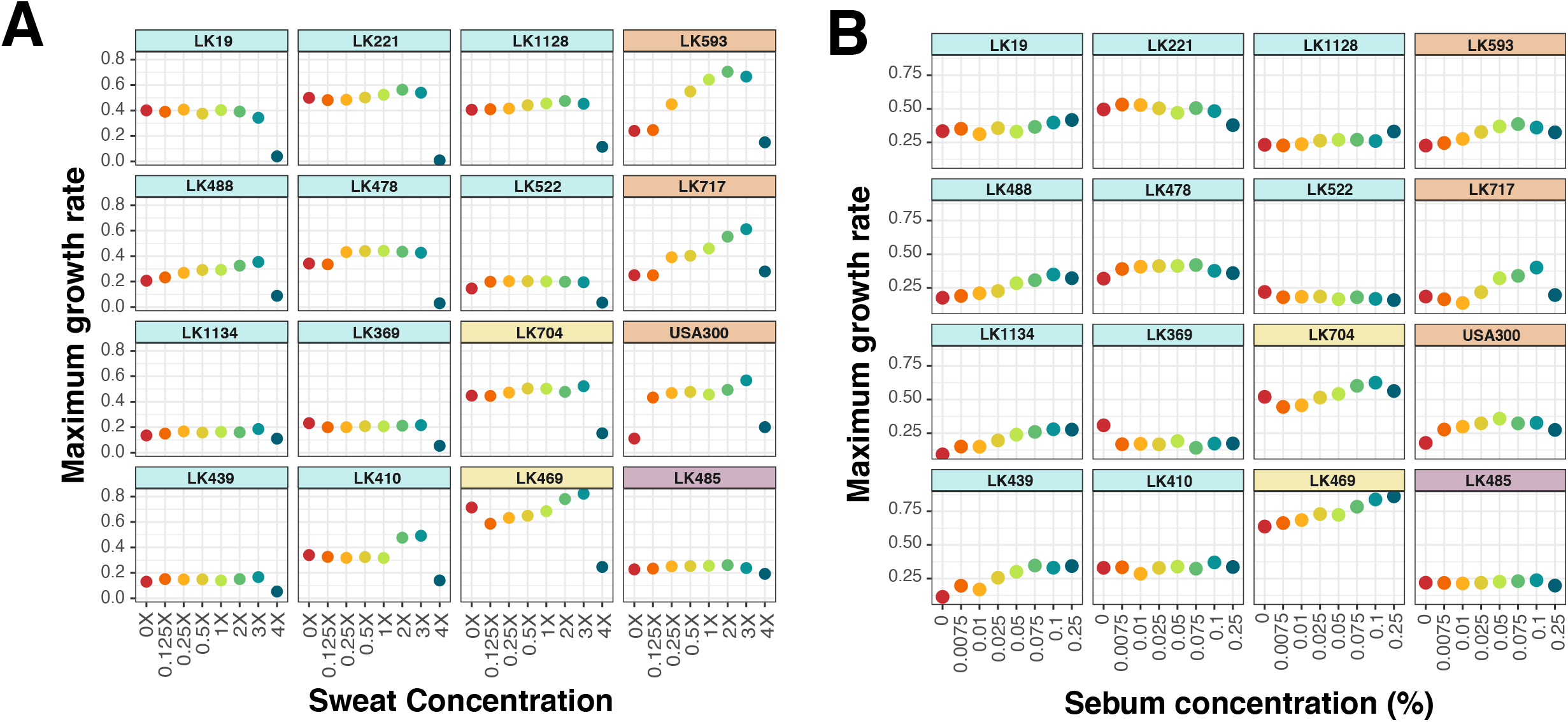
A) From the artificial sweat growth curve data, maximum growth rate was calculated for each strain and sweat concentration. B) From the artificial sebum growth curve data, maximum growth rate was calculated for each strain and sebum concentration. Strains are ordered as in Figures 2 and 3.

**Supplemental Figure 2.**
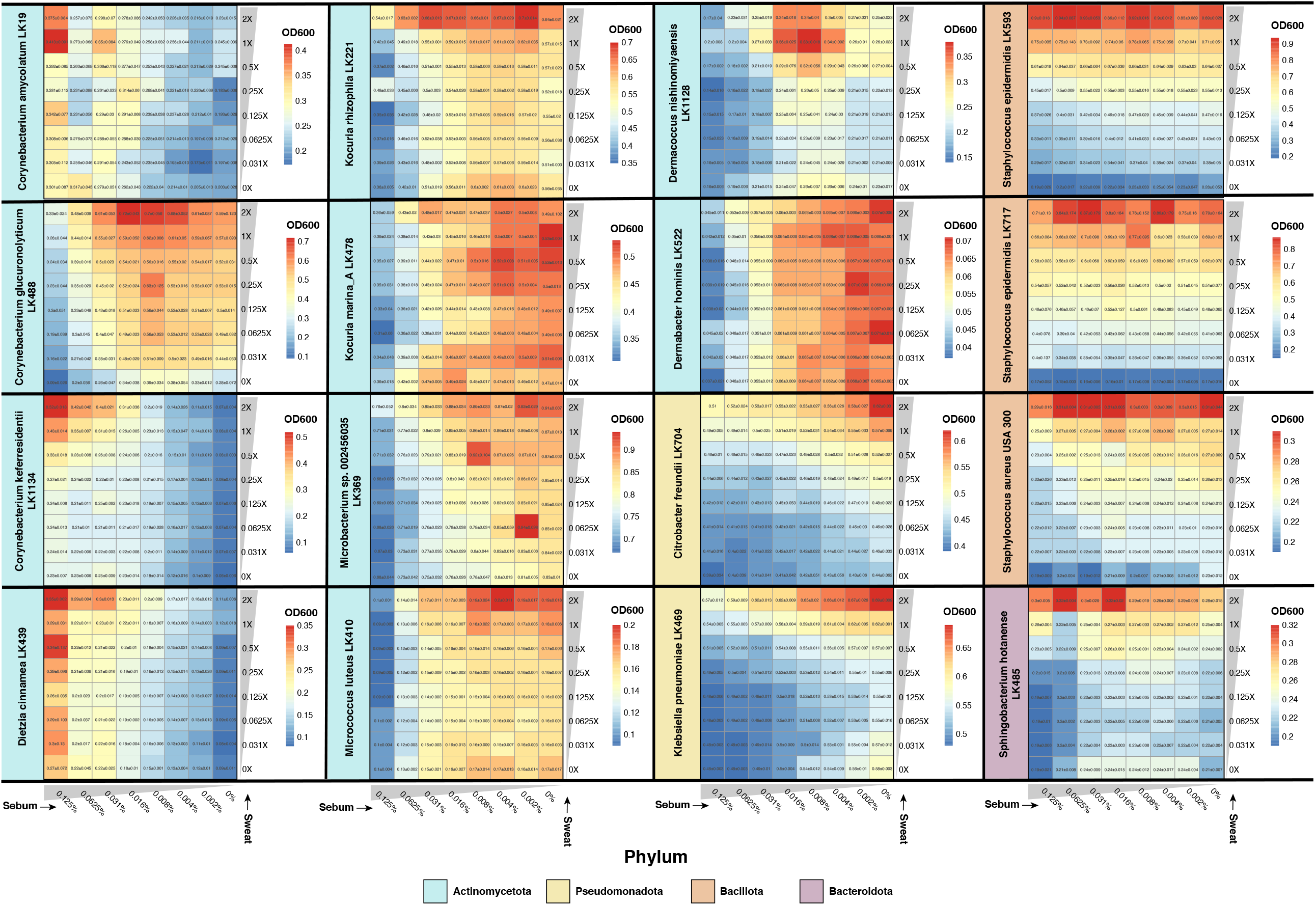
Strains were cultured in unique combinations of artificial sweat media (0, 0.031, 0.0625, 0.125, 0.25, 0.5, 1, 2X) and artificial sebum media (0, 0.002, 0.004, 0.008, 0.016, 0.031, 0.0625, 0.125%) for 18 hours, after which an endpoint OD_600_ reading was recorded. At least three replicates were collected for each strain, and OD_600_ values from uninoculated media were subtracted from experimental OD_600_ values. Average OD_600_ values of the replicates and standard deviation are indicated.

**Supplemental Figure 3.**
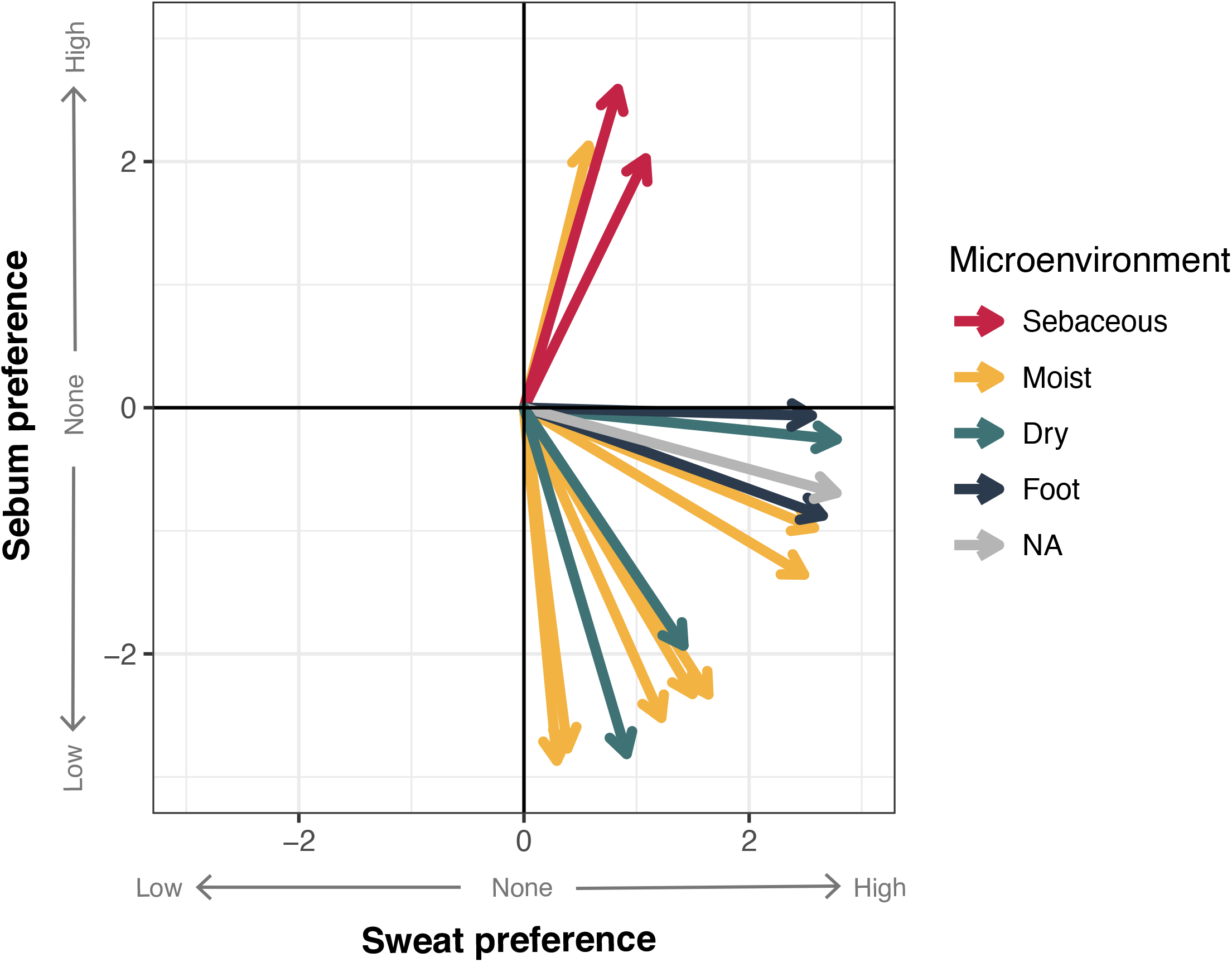
Multiple linear regression analysis was performed with the z-score normalized data from the sweat and sebum checkerboard assays. The slope coefficients for sweat concentration and sebum concentration are plotted as an (x,y) coordinate, respectively. Strains are colored by skin microenvironment.

**Supplemental Table 1.** Basal medium composition.

| Basal Medium Component | Composition (mg) | Final Concentration |
| --- | --- | --- |
| Salts |  |  |
| M9 Salts (g) | 11.28 (g) | 11.28 g/L |
| Cobalt Chloride hexahydrate (ug) | 12 (ug) | 50 nM |
| Calcium Chloride anhydrous | 11 | 0.1 mM |
| Magnesium Sulfate heptahydrate | 490 | 2 mM |
| Carbon Sources |  |  |
| Glucose (g) | 0.1<sup>a</sup>, 1.9<sup>b</sup> (g) | 2 g/L |
| Amino Acids |  |  |
| L-Arginine | 100 | 100 mg/L |
| L-Proline | 200 | 200 mg/L |
| Vitamins |  |  |
| Thiamine-HCl | 2 | 2 mg/L |
| Biotin | 2 | 2 mg/L |
| Nicotinic acid | 2 | 2 mg/L |
| Calcium pantothenate | 2 | 2 mg/L |

Values are expressed in milligrams (per 1 liter of medium) unless noted otherwise. <sup>a</sup> Added before autoclaving. <sup>b</sup> Added after autoclaving.

## METHODS

### Bacterial strains and media

Bacterial strains used in this study are part of the Kalan laboratory skin strain biobank and were isolated from the skin of healthy volunteers under an IRB approved protocol (Sandstrom *et al*. - manuscript in preparation). Strain information is listed in Supplemental Material S2. For routine culturing, strains were struck out from frozen glycerol stocks onto Brain Heart Infusion agar plates supplemented with 0.2% Tween 80 (BHITw80) and incubated at 37°C until single colonies approximately 0.5-1 mm in size formed. Between 1 and 5 colonies were inoculated into 4 mL BHITw80 broth and incubated at 37°C shaking (200 rpm) for 24 hours for use in subsequent experiments.

### Basal medium composition and preparation

A minimal medium was developed for use as a basal medium for the experiments in this study. 11.28 g/L M9 salts (Sigma-Aldrich) and 0.1 g/L glucose were prepared in aqueous solution and autoclaved at 121°C for 15 minutes. When cooled, the following media components were added at the final concentrations indicated: 0.1 mM CaCl2, 2 mM MgSO4, 50 nM CoCl2, 100 mg/L L-arginine, 200 mg/L L-proline, 2 mg/L thiamine-HCl, 2 mg/L nicotinic acid, 2 mg/L calcium pantothenate, and 2 mg/L biotin. Additional glucose solution was added to bring the final concentration to 2 g/L. As described previously, certain skin strains exhibit improved growth when a small amount of glucose is autoclaved in the presence of other medium components (21, 71). A summary of the basal medium composition is listed in Supplemental Table 1.

### Artificial sweat development

Artificial sweat was developed based on the composition of previously described synthetic sweat media (12, 14) as well as through extrapolation of sweat constituents and their concentrations as reported in the literature (18, 24, 36–45). Because a range of concentrations exist for many constituents reported in human sweat, we selected a value generally within the middle of the range of concentrations. The final artificial sweat composition used in this study can be found in Table 1. To prepare the artificial sweat, all components indicated in Table 1 were added to H_2_O to the desired concentration, followed by filter sterilization and storage at 4°C. We have had success preparing concentrated solutions of the artificial sweat up to 8X concentration.

### Artificial sebum development

Artificial sebum was developed based upon previously described artificial sebum L (27). Minor modifications were made based upon availability of individual components (replacement of spermaceti with hexadecyl palmitate, addition of myristic acid and lauric acid, exclusion of palmitoleic acid). The final composition used in this study can be found in Table 2. To prepare the artificial sebum, all components indicated in Table 2 were added to a glass bottle placed in a hot water bath. The components were heated with intermittent stirring until the solids melted to form a homogenous oil. The final product was stored at 4°C. Preparation of the sebum in aqueous solution is as follows: artificial sebum and Tween 80 were heated to an oil consistency, followed by mixing of one part artificial sebum with three parts Tween 80. The sebum/Tween 80 suspension was added to H_2_O, and the final solution was autoclaved at 121°C for 15 minutes. We have had success preparing aqueous solutions of the artificial sebum up to 0.25%.

### 16S rRNA phylogenetic tree

Ribosomal RNA sequences were identified from whole genome sequences of the 16 strains using barrnap (v0.9). 16S rRNA gene sequences were extracted, aligned with muscle (v3.8.1551), and trimmed with trimal (v1.4.rev22). Fasttree (v2.1.10) was used to infer an approximately-maximum-likelihood phylogenetic tree of the 16S rRNA sequences. The tree was rooted using the archaeum *Halobacterium salinarum*.

### Sweat and sebum growth curve assays

Eight concentrations of sweat medium (0, 0.0625, 0.125, 0.25, 0.5, 1, 2, 4X) were prepared in basal medium using a concentrated stock of 4X artificial sweat. Similarly, eight concentrations of sebum medium (0, 0.0075, 0.01, 0.025, 0.05, 0.075, 0.1, 0.25%) were prepared in basal medium using the artificial sebum/Tween 80 formulation described above. 0X sweat and 0% sebum are comprised of only basal medium, while 4X sweat is comprised of only concentrated 4X artificial sweat. The concentration of basal medium components in the final medium preparations remained constant and was adjusted for during media preparation. A liquid culture of each test strain (see **Bacterial strains and media**) was diluted to an OD_600_ of 3.0, and 10 µ L of the diluted culture was then inoculated into sweat or sebum medium in a lidded 96-well plate (CytoOne) to a final OD_600_ of 0.15 in a volume of 200 µL. We selected this OD_600_ because we have observed that many skin strains exhibit density dependent growth, where too low of an inoculum results in minimal or no growth. Assay setup is outlined in Figure 1D. The plate was incubated at 37°C shaking in an Epoch 2 microplate reader (BioTek, USA) for 36 hours, and OD_600_ measurements were recorded every 10 minutes. Three technical replicates and at least two biological replicates were collected for each strain and media condition.

For each media condition, OD_600_ technical replicates for each strain were averaged and subtracted from the average of the OD_600_ technical replicates of the uninoculated media. Averages and standard deviations were then calculated for all biological replicates of each strain and media condition. A LOESS model was fit for the biological replicates, and area under the curve (AUC) was calculated with the trapz function from the pracma R package (v2.3.8). Maximum growth rate for each strain and media concentration was calculated with the all_easylinear function from the growthrates R package (v0.8.2). Linear regression analysis was performed with min-max normalized AUC data, and the slope coefficient (β) and p-value were used to determine sweat or sebum preference. β < 0 and p-value < 0.05 indicates a low sweat or sebum preference phenotype; β > 0 and p-value < 0.05 indicates a high sweat or sebum preference phenotype; and p-value > 0.05 indicate no sweat or sebum preference.

### Sweat and sebum checkerboard assays

4X sweat medium was prepared in basal medium from a concentrated stock of 8X artificial sweat. To prepare diluted sweat media for the checkerboard assay, 4X sweat medium was added to the top row (wells 1-8) of a 96-well plate, and basal medium was added to all remaining rows (wells 1-8) (Figure 1E). Two-fold dilutions were performed down rows A-G, with the last row consisting of only basal medium. Similarly, to prepare diluted sebum media for the checkerboard assay, 0.25% sebum medium was added to the first column of another 96-well plate, and basal medium was added to the remaining columns (wells 1-8) (Figure 1E). Two-fold dilutions were performed across columns 1-8, with the last column consisting of only basal medium. 95 µL from each well of both the sweat and sebum media dilution plates were transferred to another 96-well plate, resulting in a sweat and sebum checkerboard containing 190 µL media per well.

A liquid culture of each test strain (see **Bacterial strains and media**) was diluted to an OD_600_ of 3.0, and 10 µL of the diluted culture was then inoculated into all wells of the sweat and sebum checkerboard plate. Plates were incubated at 37°C shaking (200 rpm) in a New Brunswick Innova 42 incubator (Eppendorf) for 18 hours. After incubation, cells were mixed by pipetting and OD_600_ measurements were taken in an Epoch 2 microplate reader (BioTek, USA). At least three replicates were collected for each strain.

OD_600_ values for each plate were subtracted from the average of the OD_600_ replicates of uninoculated media. Averages and standard deviations were then calculated for all replicates of each strain. Data were z-score normalized for subsequent analyses. For each strain, a multiple linear regression was performed using the formula (OD_600_ ∼ sebum concentration + sweat concentration), and the relative importance of sweat and sebum concentration in the model were determined using the calc.relimp function from the relaimpo R package (v2.2-6). Principal component analysis and hierarchical clustering (average-linkage method) were performed, followed by tanglegram analysis of the resulting dendrogram and 16S rRNA phylogeny using the tanglegram function from the dendextend R package (v1.15.2). For multiple linear regression analysis, the slope coefficients (β_Sweat_ and β_Sebum_) and p-value were used to determine sweat and sebum preference. β_Sweat_ or β_Sebum_ < 0 and p-value < 0.05 indicates a low sweat or sebum preference phenotype; β_Sweat_ or β_Sebum_ > 0 and p-value < 0.05 indicate a high sweat or sebum preference phenotype; and p-value > 0.05 indicates no sweat or sebum preference.

## REFERENCES

1. Zheng Y, Hunt RL, Villaruz AE, Fisher EL, Liu R, Liu Q, Cheung GYC, Li M, Otto M. 2022. Commensal Staphylococcus epidermidis contributes to skin barrier homeostasis by generating protective ceramides. Cell Host Microbe https://doi.org/10.1016/j.chom.2022.01.004.

2. Dwyer LR, Scharschmidt TC. 2022. Early life host-microbe interactions in skin. Cell Host Microbe 30:684–695.

3. Flowers L, Grice EA. 2020. The Skin Microbiota: Balancing Risk and Reward. Cell Host Microbe 28:190–200.

4. Swaney MH, Kalan LR. 2021. Living in Your Skin: Microbes, Molecules, and Mechanisms. Infect Immun 89.

5. Belkaid Y, Segre JA. 2014. Dialogue between skin microbiota and immunity. Science 346:954–959.

6. Byrd AL, Belkaid Y, Segre JA. 2018. The human skin microbiome. Nat Rev Microbiol 16:143–155.

7. Whiteside JA, Voss JG. 1973. Incidence and Lipolytic Activity of Propionibacterium Acnes (Corynebacterium Acnes Group I) And P. Granulosum (C. Acnes Group II) in Acne and in Normal Skin. Journal of Investigative Dermatology https://doi.org/10.1111/1523-1747.ep12724177.

8. Puhvel SM, Reisner RM. 1970. Effect of fatty acids on the growth of Corynebacterium acnes in vitro. J Invest Dermatol 54:48–52.

9. Gribbon EM, Cunliffe WJ, Holland KT. 1993. Interaction of Propionibacterium acnes with skin lipids in vitro. J Gen Microbiol 139:1745–1751.

10. Tauch A, Bischoff N, Pühler A, Kalinowski J. 2004. Comparative genomics identified two conserved DNA modules in a corynebacterial plasmid family present in clinical isolates of the opportunistic human pathogen Corynebacterium jeikeium. Plasmid 52:102–118.

11. Kwaszewska A, Sobiś-Glinkowska M, Szewczyk EM. 2014. Cohabitation--relationships of corynebacteria and staphylococci on human skin. Folia Microbiol 59:495–502.

12. Horton MV, Johnson CJ, Kernien JF, Patel TD, Lam BC, Cheong JZA, Meudt JJ, Shanmuganayagam D, Kalan LR, Nett JE. 2020. Candida auris forms high-burden biofilms in skin niche conditions and on porcine skin. mSphere 5.

13. Watson F, Keevil CW, Chewins J, Wilks SA. 2022. Artificial Human Sweat as a Novel Growth Condition for Clinically Relevant Pathogens on Hospital Surfaces. Microbiol Spectr e0213721.

14. Callewaert C, Buysschaert B, Vossen E, Fievez V, Van de Wiele T, Boon N. 2014. Artificial sweat composition to grow and sustain a mixed human axillary microbiome. J Microbiol Methods 103:6–8.

15. Otberg N, Richter H, Schaefer H, Blume-Peytavi U, Sterry W, Lademann J. 2004. Variations of hair follicle size and distribution in different body sites. J Invest Dermatol 122:14–19.

16. Taylor NA, Machado-Moreira CA. 2013. Regional variations in transepidermal water loss, eccrine sweat gland density, sweat secretion rates and electrolyte composition in resting and exercising humans. Extrem Physiol Med 2:4.

17. Smith CJ, Havenith G. 2011. Body mapping of sweating patterns in male athletes in mild exercise-induced hyperthermia. Eur J Appl Physiol 111:1391–1404.

18. Wilson M. 2005. Microbial Inhabitants of Humans: Their Ecology and Role in Health and Disease. Cambridge University Press.

19. Oh J, Byrd AL, Deming C, Conlan S, NISC Comparative Sequencing Program, Kong HH, Segre JA. 2014. Biogeography and individuality shape function in the human skin metagenome. Nature 514:59–64.

20. Oh J, Byrd AL, Park M, Kong HH, Segre JA. 2016. Temporal Stability of the Human Skin Microbiome. Cell 165:854–866.

21. Swaney MH, Sandstrom S, Kalan LR. 2021. Cobamide sharing drives skin microbiome dynamics. bioRxiv.

22. Claesen J, Spagnolo JB, Ramos SF, Kurita KL, Byrd AL, Aksenov AA, Melnik AV, Wong WR, Wang S, Hernandez RD, Donia MS, Dorrestein PC, Kong HH, Segre JA, Linington RG, Fischbach MA, Lemon KP. 2020. A Cutibacterium acnes antibiotic modulates human skin microbiota composition in hair follicles. Sci Transl Med 12.

23. Baker LB, Wolfe AS. 2020. Physiological mechanisms determining eccrine sweat composition. Eur J Appl Physiol 120:719–752.

24. Baker LB. 2019. Physiology of sweat gland function: The roles of sweating and sweat composition in human health. Temperature (Austin) 6:211–259.

25. Thody AJ, Shuster S. 1989. Control and function of sebaceous glands. Physiol Rev 69:383–416.

26. Picardo M, Ottaviani M, Camera E, Mastrofrancesco A. 2009. Sebaceous gland lipids. Dermatoendocrinol 1:68–71.

27. Lu GW, Valiveti S, Spence J, Zhuang C, Robosky L, Wade K, Love A, Hu L-Y, Pole D, Mollan M. 2009. Comparison of artificial sebum with human and hamster sebum samples. Int J Pharm 367:37–43.

28. Nielsen CK, Kjems J, Mygind T, Snabe T, Meyer RL. 2016. Effects of Tween 80 on Growth and Biofilm Formation in Laboratory Media. Front Microbiol 7:1878.

29. Nakatsuji T, Kao MC, Zhang L, Zouboulis CC, Gallo RL, Huang C-M. 2010. Sebum free fatty acids enhance the innate immune defense of human sebocytes by upregulating beta-defensin-2 expression. J Invest Dermatol 130:985–994.

30. Gerrett N, Griggs K, Redortier B, Voelcker T, Kondo N, Havenith G. 2018. Sweat from gland to skin surface: production, transport, and skin absorption. J Appl Physiol 125:459–469.

31. Ely MR, Kenefick RW, Cheuvront SN, Chinevere TD, Lacher CP, Lukaski HC, Montain SJ. 2011. Surface contamination artificially elevates initial sweat mineral concentrations. J Appl Physiol 110:1534–1540.

32. Baker LB. 2017. Sweating Rate and Sweat Sodium Concentration in Athletes: A Review of Methodology and Intra/Interindividual Variability. Sports Med 47:111–128.

33. Sprouffske K, Wagner A. 2016. Growthcurver: an R package for obtaining interpretable metrics from microbial growth curves. BMC Bioinformatics 17:172.

34. Gill SR, Fouts DE, Archer GL, Mongodin EF, Deboy RT, Ravel J, Paulsen IT, Kolonay JF, Brinkac L, Beanan M, Dodson RJ, Daugherty SC, Madupu R, Angiuoli SV, Durkin AS, Haft DH, Vamathevan J, Khouri H, Utterback T, Lee C, Dimitrov G, Jiang L, Qin H, Weidman J, Tran K, Kang K, Hance IR, Nelson KE, Fraser CM. 2005. Insights on evolution of virulence and resistance from the complete genome analysis of an early methicillin-resistant Staphylococcus aureus strain and a biofilm-producing methicillin-resistant Staphylococcus epidermidis strain. J Bacteriol 187:2426–2438.

35. Otto M. 2010. Staphylococcus colonization of the skin and antimicrobial peptides. Expert Rev Dermatol 5:183–195.

36. Mark H, Harding CR. 2013. Amino acid composition, including key derivatives of eccrine sweat: potential biomarkers of certain atopic skin conditions. Int J Cosmet Sci 35:163–168.

37. Robinson S, Robinson AH. 1954. Chemical composition of sweat. Physiol Rev 34:202–220.

38. Thurmon FM, Ottenstein B. 1952. Studies on the chemistry of human perspiration with especial reference to its lactic acid content. J Invest Dermatol 18:333–339.

39. Montain SJ, Cheuvront SN, Lukaski HC. 2007. Sweat mineral-element responses during 7 h of exercise-heat stress. Int J Sport Nutr Exerc Metab 17:574–582.

40. Patterson MJ, Galloway SD, Nimmo MA. 2000. Variations in regional sweat composition in normal human males. Exp Physiol 85:869–875.

41. Dunstan RH, Hugh Dunstan R, Sparkes DL, Dascombe BJ, Macdonald MM, Evans CA, Stevens CJ, Crompton MJ, Gottfries J, Franks J, Murphy G, Wood R, Roberts TK. 2016. Sweat Facilitated Amino Acid Losses in Male Athletes during Exercise at 32-34°C. PLOS ONE https://doi.org/10.1371/journal.pone.0167844.

42. Harvey CJ, LeBouf RF, Stefaniak AB. 2010. Formulation and stability of a novel artificial human sweat under conditions of storage and use. Toxicol In Vitro 24:1790–1796.

43. Huang C-T, Chen M-L, Huang L-L, Mao I-F. 2002. Uric acid and urea in human sweat. Chin J Physiol 45:109–115.

44. Mitchell HH, Hamilton TS. 1949. The dermal excretion under controlled environmental conditions of nitrogen and minerals in human subjects, with particular reference to calcium and iron. J Biol Chem 178:345–361.

45. Lee H, Song C, Hong YS, Kim MS, Cho HR, Kang T, Shin K, Choi SH, Hyeon T, Kim D-H. 2017. Wearable/disposable sweat-based glucose monitoring device with multistage transdermal drug delivery module. Sci Adv 3:e1601314.

46. Lam TH, Verzotto D, Brahma P, Ng AHQ, Hu P, Schnell D, Tiesman J, Kong R, Ton TMU, Li J, Ong M, Lu Y, Swaile D, Liu P, Liu J, Nagarajan N. 2018. Understanding the microbial basis of body odor in pre-pubescent children and teenagers. Microbiome 6:213.

47. Scharschmidt TC, Fischbach MA. 2013. What Lives On Our Skin: Ecology, Genomics and Therapeutic Opportunities Of the Skin Microbiome. Drug Discov Today Dis Mech 10.

48. Kunin CM, Rudy J. 1991. Effect of NaCl-induced osmotic stress on intracellular concentrations of glycine betaine and potasxsium in Escherichia coli, Enterococcus faecalis, and staphylococci. J Lab Clin Med 118:217–224.

49. Onyango LA, Alreshidi MM. 2018. Adaptive Metabolism in Staphylococci: Survival and Persistence in Environmental and Clinical Settings. J Pathog 2018:1092632.

50. Rahman SS, Siddique R, Tabassum N. 2017. Isolation and identification of halotolerant soil bacteria from coastal Patenga area. BMC Res Notes 10:531.

51. Ben-Dov E, Ben Yosef DZ, Pavlov V, Kushmaro A. 2009. Corynebacterium maris sp. nov., a marine bacterium isolated from the mucus of the coral Fungia granulosa. Int J Syst Evol Microbiol 59:2458–2463.

52. Rückert C, Albersmeier A, Al-Dilaimi A, Niehaus K, Szczepanowski R, Kalinowski J. 2012. Genome sequence of the halotolerant bacterium Corynebacterium halotolerans type strain YIM 70093(T) (= DSM 44683(T)). Stand Genomic Sci 7:284–293.

53. Irlinger F, Helinck S, Jany JL. 2017. Chapter 11 - Secondary and Adjunct Cultures, p. 273– 300. In McSweeney, PLH, Fox, PF, Cotter, PD, Everett, DW (eds.), Cheese (Fourth Edition). Academic Press, San Diego.

54. Bharti N, Pandey SS, Barnawal D, Patel VK, Kalra A. 2016. Plant growth promoting rhizobacteria Dietzia natronolimnaea modulates the expression of stress responsive genes providing protection of wheat from salinity stress. Sci Rep 6:34768.

55. Stackebrandt E, Koch C, Gvozdiak O, Schumann P. 1995. Taxonomic dissection of the genus Micrococcus: Kocuria gen. nov., Nesterenkonia gen. nov., Kytococcus gen. nov., Dermacoccus gen. nov., and Micrococcus Cohn 1872 gen. emend. Int J Syst Bacteriol 45:682–692.

56. Novitsky TJ, Kushner DJ. 1975. Influence of temperature and salt concentration on the growth of a facultatively halophilic “Micrococcus” sp. Can J Microbiol 21:107–110.

57. Yang Z-W, Salam N, Mohany M, Chinnathambi A, Alharbi SA, Xiao M, Hozzein WN, Li W-J. 2018. Microbacterium album sp. nov. and Microbacterium deserti sp. nov., two halotolerant actinobacteria isolated from desert soil. Int J Syst Evol Microbiol 68:217–222.

58. Orji OU, Awoke JN, Aja PM, Aloke C, Obasi OD, Alum EU, Udu-Ibiam OE, Oka GO. 2021. Halotolerant and metalotolerant bacteria strains with heavy metals biorestoration possibilities isolated from Uburu Salt Lake, Southeastern, Nigeria. Heliyon 7:e07512.

59. Tóth Á, Bata-Vidács I, Kosztik J, Máté R, Kutasi J, Tóth E, Bóka K, Táncsics A, Nagy I, Kovács G, Kukolya J. 2021. Sphingobacterium pedocola sp. nov. a novel halotolerant bacterium isolated from agricultural soil. Antonie Van Leeuwenhoek 114:1575–1584.

60. Tauch A, Kaiser O, Hain T, Goesmann A, Weisshaar B, Albersmeier A, Bekel T, Bischoff N, Brune I, Chakraborty T, Kalinowski J, Meyer F, Rupp O, Schneiker S, Viehoever P Pühler 2005. Complete genome sequence and analysis of the multiresistant nosocomial pathogen Corynebacterium jeikeium K411, a lipid-requiring bacterium of the human skin flora. J Bacteriol 187:4671–4682.

61. Salamzade R, Alex Cheong JZ, Sandstrom S, Swaney MH, Starr NL, Singh AM, Kalan LR. 2022. lsaBGC provides a comprehensive framework for evolutionary analysis of biosynthetic gene clusters within focal taxa. bioRxiv.

62. Wille JJ, Kydonieus A. 2003. Palmitoleic Acid Isomer (C16:1Δ6) in Human Skin Sebum Is Effective against Gram-Positive Bacteria. Skin Pharmacol Physiol 16:176–187.

63. Drake DR, Brogden KA, Dawson DV, Wertz PW. 2008. Thematic review series: skin lipids. Antimicrobial lipids at the skin surface. J Lipid Res 49:4–11.

64. Chamberlain NR, Brueggemann SA. 1997. Characterisation and expression of fatty acid modifying enzyme produced by Staphylococcus epidermidis. J Med Microbiol 46:693–697.

65. Brüggemann H, Henne A, Hoster F, Liesegang H, Wiezer A, Strittmatter A, Hujer S, Dürre P, Gottschalk G. 2004. The complete genome sequence of Propionibacterium acnes, a commensal of human skin. Science 305:671–673.

66. Miskin JE, Farrell AM, Cunliffe WJ, Holland KT. 1997. Propionibacterium acnes, a resident of lipid-rich human skin, produces a 33 kDa extracellular lipase encoded by gehA. Microbiology 143 (Pt 5):1745–1755.

67. Brüggemann H, Salar-Vidal L, Gollnick HPM, Lood R. 2021. A Janus-Faced Bacterium: Host-Beneficial and -Detrimental Roles of Cutibacterium acnes. Front Microbiol 12:673845.

68. Rieg S, Seeber S, Steffen H, Humeny A, Kalbacher H, Stevanovic S, Kimura A, Garbe C, Schittek B. 2006. Generation of multiple stable dermcidin-derived antimicrobial peptides in sweat of different body sites. J Invest Dermatol 126:354–365.

69. Rieg S, Garbe C, Sauer B, Kalbacher H, Schittek B. 2004. Dermcidin is constitutively produced by eccrine sweat glands and is not induced in epidermal cells under inflammatory skin conditions. Br J Dermatol 151:534–539.

70. Murakami M, Ohtake T, Dorschner RA, Schittek B, Garbe C, Gallo RL. 2002. Cathelicidin anti-microbial peptide expression in sweat, an innate defense system for the skin. J Invest Dermatol 119:1090–1095.

71. Liebl W, Klamer R, Schleifer K-H. 1989. Requirement of chelating compounds for the growth of Corynebacterium glutamicum in synthetic media. Appl Microbiol Biotechnol 32:205–210.

